# Feeding Ecology and Behavioral Adaptations Shape Injury Patterns in Central European Ants

**DOI:** 10.64898/2026.02.09.704771

**Authors:** Melvin Kenneth Opolka, Alina Koeters, Erik Thomas Frank

## Abstract

Injuries are common in animals and represent a major threat to individual survival. They can result from inter- or intraspecific conflict, predation, or pugnacious prey. Despite their potential ecological and evolutionary importance, injury patterns remain poorly documented in animal populations. To test whether a species’ feeding ecology or habitat can predict injury patterns, we quantified injury rates and affected body regions among native ant species collected from different habitats in Bavaria, Germany. Specimens were sampled using pitfall traps, which proved to be an efficient method for injury assessment.

Injury rates varied substantially among species and genera, ranging from 0% to 38%. Predatory ant species exhibited higher frequencies of leg injuries, whereas omnivorous species were more frequently injured at the antennae. The distribution of injuries likely reflects both foraging ecology and species-specific wound care behaviors, with a high frequency of trochanter injuries potentially indicating prior amputation events to cope with infected leg injuries, as observed in *Lasius alienus*.

Our findings demonstrate that injury propensity and distribution are shaped by feeding habits and behavioral adaptations, providing comparative evidence that the costs and management of injuries vary systematically among ant species. Our study thus highlights injuries as a measurable axis of selection that may have contributed to the emergence of wound care and other forms of social immunity in ants.

## 1. Introduction

Injuries are widespread in animals and have severe consequences for their fitness. Causes of injuries can include predation and fights with conspecifics over territory, food, or mating partners (Bowerman et al., 2010; de Ramirez et al., 2012; Huntingford & Turner, 1987). Injuries can lead to the loss of bodily fluids or impair an individual’s movement while searching for food or mating partners (Beamish & O’Riain, 2014; Krause et al., 2017). Ultimately, injuries also diminish both survival and reproductive success by increasing the risk of lethal infections (Carey et al., 2009; Frank et al., 2023, 2024; Moroń et al., 2012). Yet it is poorly understood how frequently injuries occur in nature and what adaptations animals have developed to mitigate their negative impacts.

Injury rates and related adaptations have mostly been studied at the level of individual species (Pusceddu et al., 2025; Subasi et al., 2024). Social animals are particularly noteworthy in this context, as they often exhibit behavioral responses to injuries. Primates are known to swallow unchewed leaves to expel intestinal parasites such as nematodes, as well as to alleviate inflammation and associated symptoms (Huffman, 1997; Huffman & Seifu, 1989; Wrangham, 1995). In addition, plant-based or inorganic substances are frequently applied externally to the fur or on wounds for their antiparasitic, antibacterial, or anti-inflammatory effects (Baker, 1996; Huffman, 1997; Laumer et al., 2024).

The best-studied examples for injury care are in social insects, particularly in ants. Workers in the termite-hunting ant *Megaponera analis* not only rescue their wounded nestmates during raids on termites by carrying them back to the nest (Frank et al., 2017) but they can also identify infected wounds and treat them with antimicrobial compounds produced in their metapleural glands, thereby drastically reducing the mortality rate of the treated nestmates (Frank et al., 2018, 2023). Wound care behaviors have also been observed in various other ant species, including generalist *Camponotus* ants (Frank et al., 2024; Fujimoto et al., 2025) which amputate the limbs of injured individuals to stop pathogen spread, and short-lived *Cataglyphis nodus* foragers, which survive for only about six days as foragers (Beydizada et al., 2024). In *Formica cinerea*, injuries reduce survival, and injured workers showed increased rescue behaviour toward intact nestmates (Turza & Miler, 2023). Even in *Eciton burchellii*, where colonies can reach over 1 million ants, a 2-tiered wound care system has evolved, including first aid at the hunting site and antimicrobial wound care with the metapleural gland inside the bivouac (Lagos-Oviedo et al., 2025). This diversity in colony size, lifespan, natural history, and wound care behaviors across species highlights the remarkable adaptability and ubiquity of wound care strategies. Theoretical models further underline their importance in increasing overall colony fitness under high injury rates (Lagos-Oviedo et al., 2025). Injuries therefore appear to be an overlooked factor affecting the ecological success of ants (Pull, 2024).

Yet while the importance of injury management in ants has been well studied behaviorally, its ecological importance, the causes and likelihood of injuries remain poorly understood. In social insects, injuries most likely occur while foraging outside the nest, by predators, prey or competitors for food or nesting sites (Boulay et al., 2007; Frank et al., 2017; Lima, 2002; Mukherjee & Heithaus, 2013; Nonacs & Dill, 1988). The life traits of ant species, including their habitat preferences and dietary habits, are therefore crucial determinants of their susceptibility to injuries.

The objective of this study is therefore to analyze injury rates among Central European ant species in Bavaria, Germany, to identify potential factors driving these injury rates, such as habitat preference, natural history or dietary habits. We hypothesize that predatory ant species will exhibit higher rates of injury compared to omnivorous ant species. Furthermore, we expect the distribution of injury types (leg or antenna) to differ between species depending on their natural history and the most likely injury sources.

## 2. Materials and Methods

### 2.1 Study region and study sites

The study was conducted in May 2022 at 44 sites throughout Bavaria (Germany) covering a region of approximately 45,000 km^2^ (Figure 1). Most plots were on Keuper formation, a fertile soil type with excellent water retention properties (Böse et al., 2018). The average temperature in the region for the period from 2003 to 2023 varied between 8.4 °C and 11.2 °C and precipitation ranged between 413 mm and 792 mm for the same period (Kaspar et al., 2013). The sites used in this study can be divided into four habitat types: grassland, forest edge, slope and wooded strip. In total, we had 10 plots with grasslands, 17 plots with forest edges, 9 plots with slopes and 8 plots with wood strips. Grasslands were defined as open landscapes characterized by poor soils and low-growing vegetation (Fig. S1E). The forest edge was the transitional area between a forest and adjacent open habitats such as meadows or fields (Fig. S1C&G). A slope was defined as the terrain along an incline where the land either rises or falls (Fig. S1A&F). The habitat wood strip was a narrow section of wooded land that often runs along fields or pathways (Fig. S1B&D).

**Figure 1.**
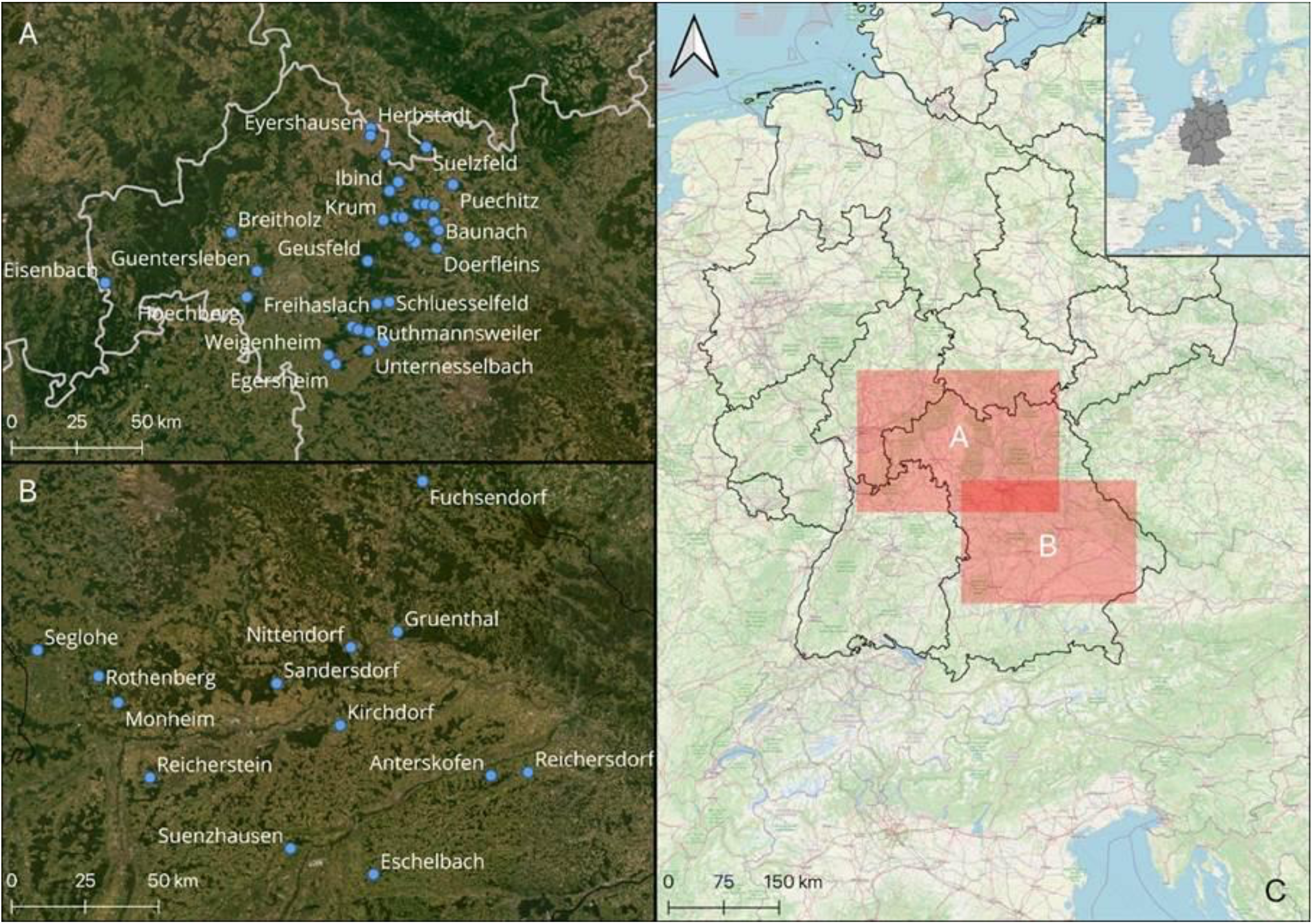
Study area in Bavaria, Germany with blue dots marking all plots used in this study. (A) Thirty-one plots situated in the upper north of Bavaria (study region A). (B) Thirteen plots are located further south in central Bavaria (study region B). (C) Map of Germany with red areas highlighting the two study regions in Bavaria where the plots were located. The top right insert includes an overview of Germany (in grey) and its neighboring countries. Map created with QGIS 3.32.3-Lima using ESRI “Satellite” basemap (ESRI, 2019).

### 2.2 Ant community sampling

For each plot, five sampling points were created in a line with one meter between each other. On each sampling point, a pitfall trap was placed, buried at soil level, and filled with a 1:2 propylene glycol-water solution, with a perfume-free detergent. The pitfall traps had a diameter of 82 mm and were recollected after two weeks. This resulted in 220 sampling points (five traps per plot, across 44 plots).

### 2.3 Ant identification and quantification of the injury rate

Pitfall trap samples were transported to the Department of Animal Ecology and Tropical Biology (Zoology III) at Julius-Maximilians University of Würzburg. All ants were transferred to 70 % ethanol for later identification. Identification of the ants was done to species level, based on keys provided by Lebas et al. 2019 and Seifert, 2007 (Table S1). Each species was then assigned to one of three types of feeding ecology: omnivore, predator, and trophobiont (Table S2). All samples were stored at the University of Würzburg (Germany). The injury rate was quantified by using a binocular (Nikon, SMZ645) with 6.7-50x magnification, to verify the state of all extremities of every ant. An injury was defined as either a complete or partially severed extremity (Figure S2). During sample processing, we took great care to avoid breaking extremities. Only genuine injuries were counted; any detached extremities found in the samples were identified as processing artefacts and excluded from the injury counts. For each observed injury, we recorded the location (front, middle, hind leg or antenna), side (left or right), and the specific injury site (funiculus, scapus, trochanter, femur, tibia or tarsus). Injury rate was then calculated as the number of injuries (divided by the total number of ants per species and pitfall.

### 2.4 Hand-collected injury rate

To verify that our sampling method with pitfalls is valid for identifying injuries in ants, we compared our injury rate of ants sampled with pitfalls with that of ants that have been collected by hand. In June 2024, approximately 100 foraging workers from 10 different colonies of *Lasius niger* were hand-collected and stored in 70 % ethanol. After the hand collection, identical pitfall traps, constructed as before, were placed next to the same 10 *L. niger* colonies for 2 weeks, and the injury rate of the trapped workers was quantified as described above.

### 2.5 Statistical analysis

For statistical analyses and graphical illustration, we utilized the R software (version 4.4.2) in combination with the RStudio interface (version 2024.12.0+467) and the *ggplot2* package (version 3.4.4) (Wickham & Chang, 2016). Our dataset contained many species that were only found a few times. To reduce the influence of very small samples, only records with at least five individuals were retained. To increase the robustness of our injury rate estimates, we excluded all ant species for which fewer than 200 individuals were collected, and which occurred in less than 6 plots. These criteria reduced the number of species analyzed from 34 to 8. Data were first tested for normality using the Shapiro-Wilk test and visualized with Q-Q plots. Normality was not met for the variable’s *habitat, ant species, genus, body side*, and *feeding ecology*, so Kruskal-Wallis tests were performed to assess differences among groups. To check for differences between injuries occurring on the left and right sides of ants, a Wilcoxon rank sum test was conducted. To analyze factors influencing injury rates, we fitted linear mixed-effects models (LMMs) using the *lme4* package (version 1.1) (Bates et al., 2023). The primary model assessed the effects of feeding ecology and habitat on injury rate, including location as a random effect to account for spatial variation. Model selection was based on Akaike’s Information Criterion (AIC). Furthermore, we applied an LMM assessing the influence of body part and feeding ecology on injury rates, with the collection site (plot) as a random effect. Model assumptions were verified by checking residual distributions using the package *DHARMa* version 0.4.7 (Hartig, 2016). Post hoc comparisons were conducted using Dunn’s test with Bonferroni correction for multiple comparisons using the FSA package version 0.6.9 (Ogle et al., 2025). We conducted a Wilcoxon rank sum test to examine how the sampling method affected injury rates.

## 3. Results

The 220 pitfall traps, distributed across the 44 plots, collected 9475 ants from 3 subfamilies, belonging to 12 genera, for a total of 34 ant species (Figure 2). The mean number of ants caught in each plot was 214.6 (± 248.9 SD, n = 44). Overall, 1137 of the 9475 ants (12%) were injured at least once. The most common ant genera in our study were *Formica, Lasius* and *Myrmica*, with *Myrmica ruginodis* being by far the most common ant species, occurring at 42 of 44 sampling sites (Table S1). While the slope habitat had a slightly higher Shannon diversity index (H’) than the other habitats (slope: H’ = 2.16; woodstrip: H’ = 2.11; grassland: H’ = 2.05; forest edge: H’ = 2.00), these differences were not significant (χ^2^ = 3, df = 3, p = 0.39).

**Figure 2.**
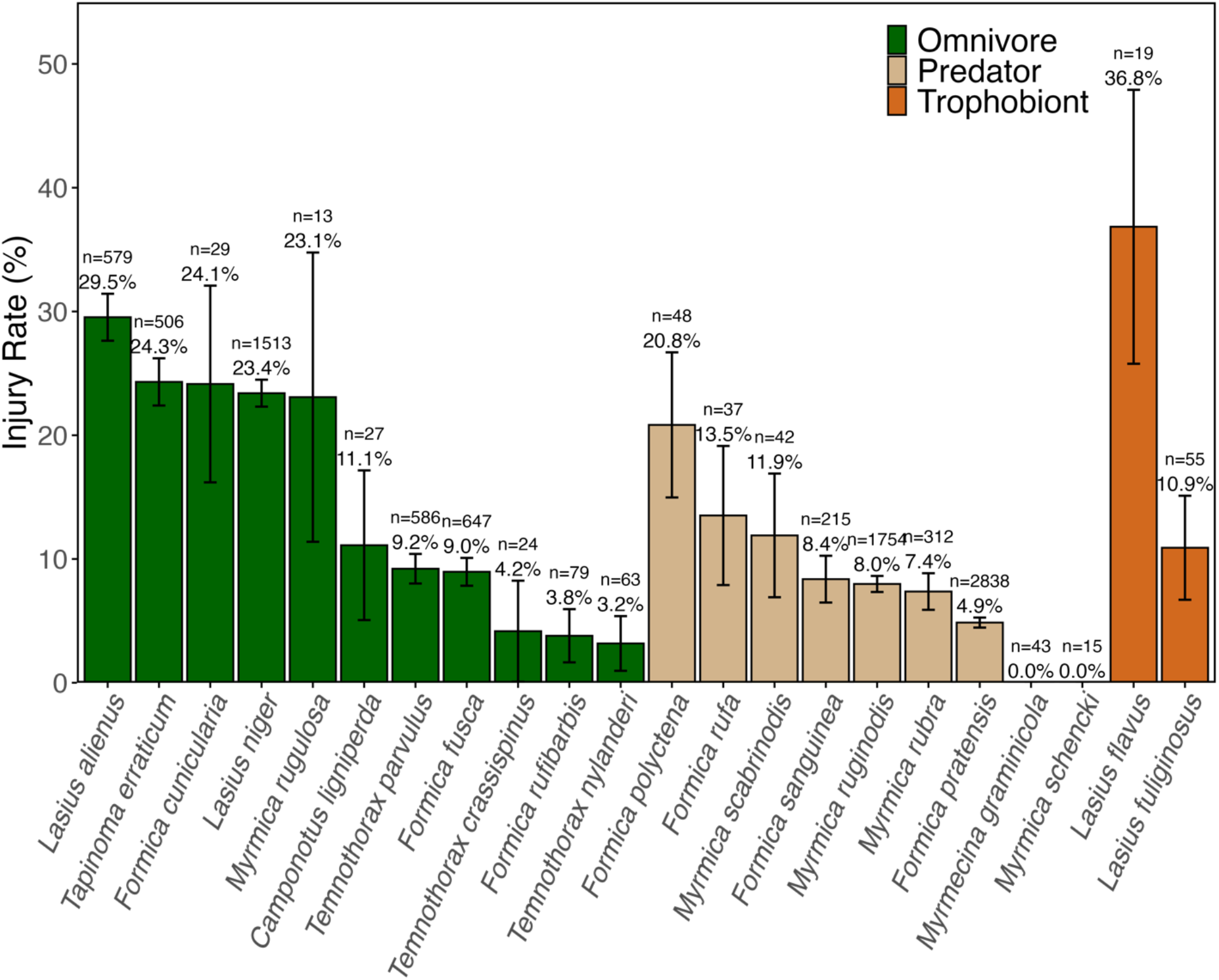
All ants found in the 44 plots, along with their injury rates and sample sizes for each species, given above the bars. Error lines indicate the binomial standard error of injury rates. The colors of the bars indicate feeding ecology: green for omnivorous species, brown for predatory species, and orange for trophobiont species.

After sorting out the rare species, we counted 983 ants with injuries, for a global injury rate of 15.8 % (n = 8205 individuals) for the remaining eight most common ant species studied. A total of 747 ants (76%) had only one injury, 179 (18%) had two injuries, and 58 (6%) were injured more than twice. The average injury rate per plot was 16.5 % for these eight species (± 8.7 SD, n = 44).

The injury rate varied significantly across species. A high injury rate was observed in *Lasius alienus* (23.1 %), *Lasius niger* (25 %) and *Tapinoma erraticum* (21.4 %). In contrast, the wood ant *Formica pratensis* (4 %) and *Temnothorax parvulus* (10.1 %) showed a low injury rate (Figure 3A).

**Figure 3.**
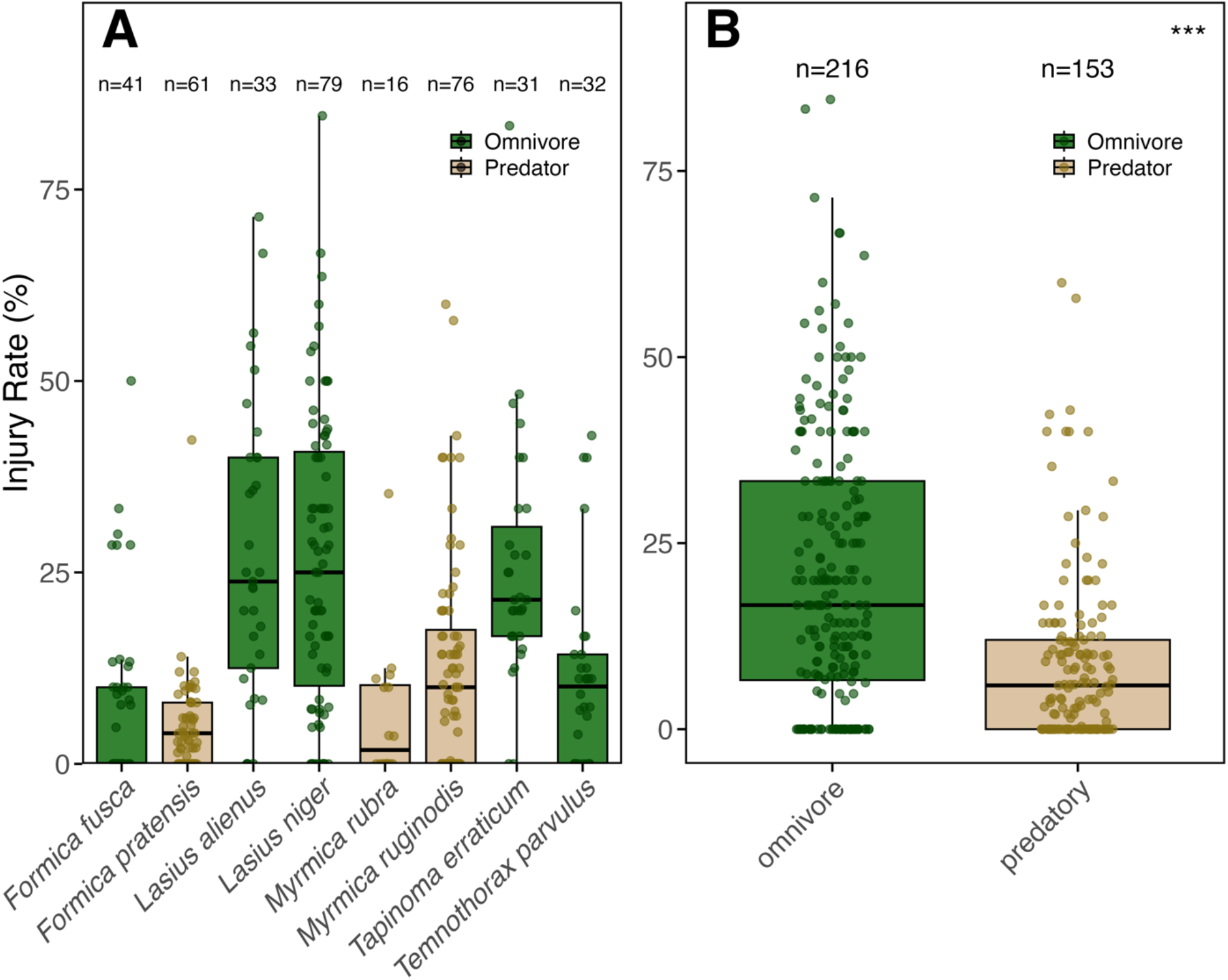
Filtered dataset showing the eight species with at least 200 sampled individuals and occurrences in more than five plots. Sample sizes are shown above the boxes. Colors indicate feeding ecology: green for omnivorous and brown for predatory species. A) Injury rates for the eight most common ant species, with each point representing one pitfall trap. B) Pooled injury rates for omnivorous and predatory species shown in A. Omnivorous species exhibit significantly higher injury rates than predatory species (LMM, Estimate = -10.84 ± 3.87, t = -2.81, p = 0.006). Each point represents the injury rate per species and pitfall.

Injuries were equally likely to occur on the left or right side of the body (Wilcoxon rank sum test: W = 68693; p = 0.83). Model comparison based on AIC values for the causes of injury rates (e.g. feeding ecology and habitat) indicated that the reduced model, which included only feeding ecology as a fixed effect, provided the best fit, while habitat and its interaction with feeding ecology showed no significant influence on injury rates (all p > 0.1; Figure 4, Table S3). Predatory species exhibited lower injury rates (9 ± 11.6%, n = 4698 ants) compared to omnivorous species (20.7 ± 18.4 %, n = 3507 ants; LMM: Estimate = -10.84 ± 3.87, t = -2.81, p = 0.006; Figure 3B). Further analyses incorporating genus and species as additional predictors did not improve model fit and were therefore not retained in the final model.

**Figure 4.**
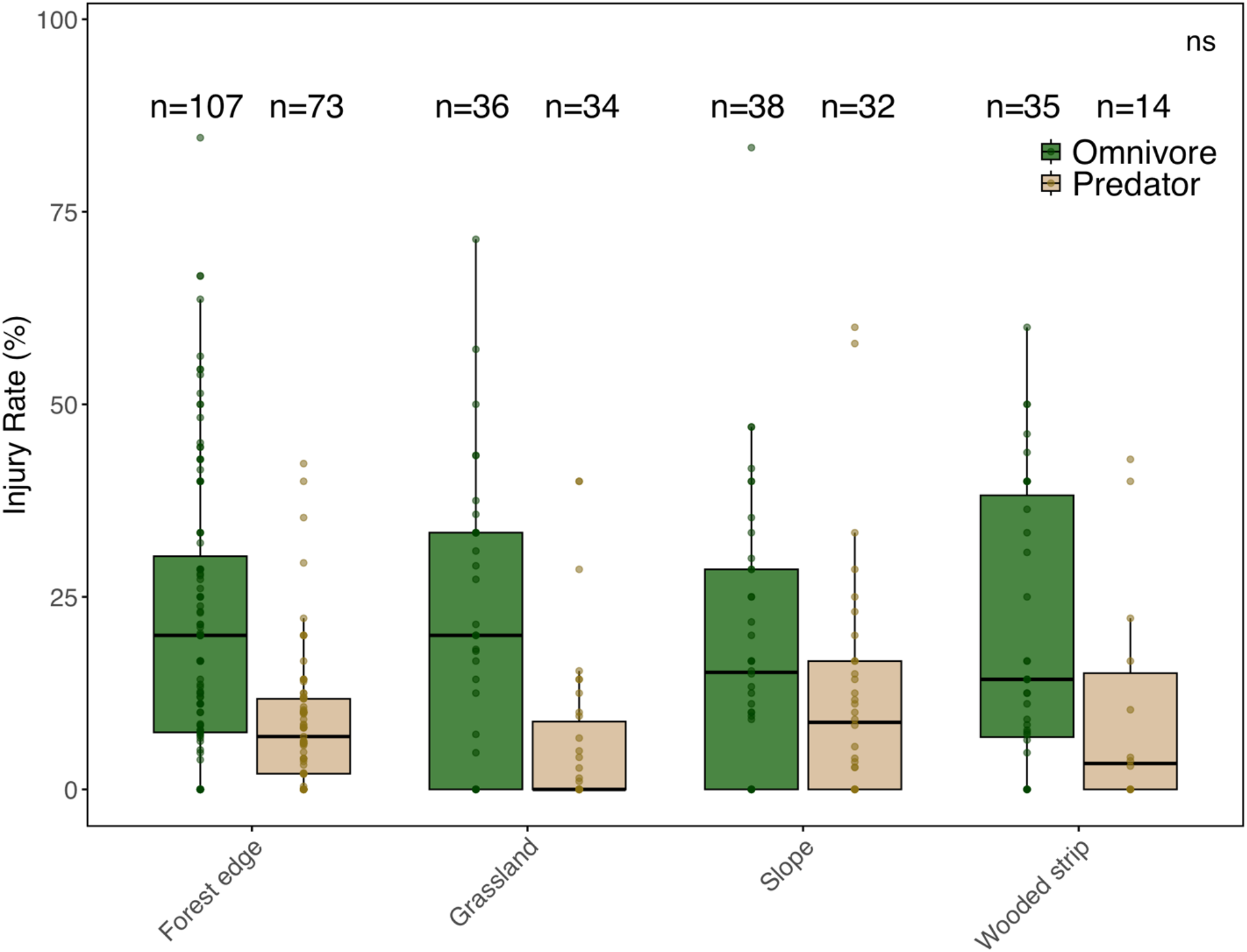
Filtered dataset showing injury rates in our four sampled habitat types. Each data point represents the injury rate in a single pitfall for each species. Sample size is indicated above the boxes, and colors indicate feeding ecology: omnivorous in green and brown for predatory species.

Injury rates differed significantly across body parts and between feeding ecology strategies, with a significant interaction between both factors (LMM: feeding ecology × body part, all p < 0.001; Table S4). Omnivorous ants exhibited higher rates of antennal injuries, while predatory ants showed relatively more injuries on the legs. Post-hoc comparisons confirmed significantly higher injury rates on the antennae than at any leg pair when analyzing the pooled dataset of omnivorous and predatory species (Antenna vs. Front Legs: t = 13.77, p < 0.0001; Antenna vs. Middle Legs: t = 15.17, p < 0.0001; Antenna vs. Hind Legs: t = 14.52, p < 0.0001; Tukey-adjusted pairwise comparisons; Figure 5A), while no significant differences were found in the injury rates across the leg pairs themselves (all p > 0.05). Nevertheless, the injury distribution varied significantly among ant species (Chi^2^, χ^2^ = 229.69, df = 21, p < 0.0001), indicating that some species experienced disproportionate injury rates in specific body regions. Notably, this effect was particularly evident when examining injuries on the trochanter and femur. Some species displayed a relatively equal distribution of injuries between the two segments, whereas others exhibited much higher injury rates concentrated on either the trochanter or the femur (Figure 5B).

**Figure 5.**
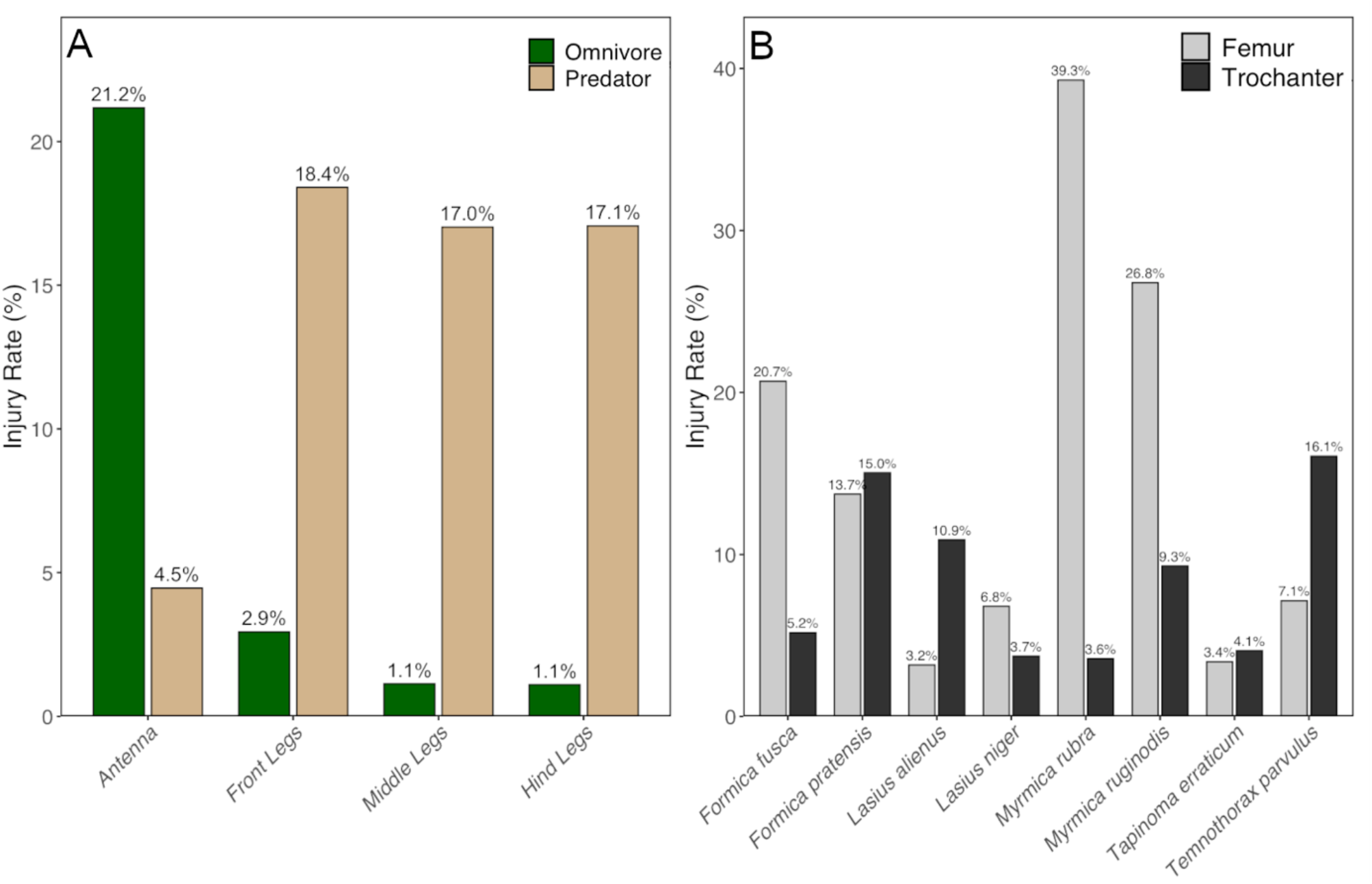
Filtered dataset showing mean injury rates (%) at A) different body part regions of omnivorous and predatory ant species. The total number of injuries studied was n = 1202 in 983 injured ants, as some ants had multiple injuries. The bars are divided into feeding ecologies of ant species, with green for omnivorous species (n = 702) and brown for predatory species (n = 281). B) Average Injury rates on the trochanter (black) and femur (grey) for the 8 most abundant ant species in our study (n = 240 injuries).

Ultimately, to test whether injuries occurred because of the pitfall traps, we compared injury rates among *Lasius niger* workers collected from 10 pitfall traps with those collected by hand at the same location. There was no significant difference between the 302 workers captured with the pitfall trap (injury rate 34 ± 16.1 %, n = 10) compared to the hand-collected 1004 workers (21.8 ± 7.9 %, n = 10; t-test: t = −2.15, df = 13.1, p = 0.051; Figure S3). Even when considering the data on *Lasius niger* from the original data set, there was no significant difference between the methods (Kruskal–Wallis test: χ^2^ = 2.99, df = 2, p = 0.224; Figure S3).

## 4. Discussion

This is the first study to quantify injury rates of ant populations across different habitats. We show how simple pitfall traps can be used to quantify these injury rates, a cost-effective and time-efficient approach. Injuries occurred across all species but followed distinct patterns related to species feeding ecologies: predatory species had a higher rate of leg injuries, while omnivorous ants showed significantly more antenna injuries.

Differences in injury rates and locations in omnivorous ants could be explained by ritualized confrontations that typically involve head-to-head interactions (Lima Vieira et al., 2024; Wilgenburg et al., 2005). During these close-range displays, the antennae are directly exposed and therefore particularly prone to injury. Omnivorous ant species, which do not actively hunt prey, may thus have a lower risk of leg injuries due to the lack of high-risk interactions with prey. However, their dependence on resources such as honeydew brings them into frequent contact with both conspecifics and other ant species (McGlynn, 2000). As a result, more frequent ritualized or direct confrontations may help explain the elevated antennal injury rates observed in these species. However, such behaviors are not exclusive to non-predatory ants and are also reported in predatory lineages such as army ants (Baudier & Pavlic, 2020; Yao, 2014).

While differences in injury location can be explained by the type of interactions typical for omnivorous species, their overall higher injury rates were unexpected. Contrary to our hypothesis, omnivorous ants showed higher injury frequencies than predatory species, which were expected to be more frequently or severely injured due to their aggressive prey encounters (Frank et al., 2017; Lagos-Oviedo et al., 2025). A possible explanation is that omnivorous species in our study are generally smaller and less well armed, (Seifert, 2007), making them more vulnerable during interspecific conflicts. Predatory species, in contrast, usually have a thicker cuticle that makes injuries less likely, whereas many Formicinae, like the omnivores in our study, possess a notably thin cuticle and are therefore more prone to mechanical damage (Peeters et al., 2017). As a result, predatory species may exhibit fewer visible injuries overall, while in Formicinae, even a single aggressive encounter could carry a higher risk of losing an extremity.

Injury rates were the same across all four habitat types, a pattern that was also consistent across both feeding ecology groups. This suggests that species specific behaviors, rather than environmental conditions, play a predominant role in shaping injury frequencies. This pattern is in line with recent work proposing that injury risk and lethality, largely arising from biotic interactions such as predator prey interactions or competition, may serve as key drivers underlying the evolution of care behaviors in ants, rather than abiotic or structural habitat factors (Frank et al., 2017; Lagos-Oviedo et al., 2025; Rennolds & Bely, 2023). This interpretation is consistent with the absence of significant differences in Shannon diversity among habitats, although this does not allow conclusions about causal relationships with injury frequencies.

The injury rate and its distribution may also offer indications on the presence of wound care behaviors. A high injury rate among foragers not only suggests frequent exposure to injury risks, but also implies that at least some individuals successfully recover from injuries, potentially through wound care, since injuries observed in the field are likely from previous foraging trips. Recent theoretical models propose that high injury rates favor the evolution of wound care behaviors, independent of colony size, making injury rates the main predictor for the presence of wound care behaviors(Frank et al., 2017, 2018; Lagos-Oviedo et al., 2025). Our study adds to this by showing how such patterns may be shaped by ecological factors such as habitat or diet. Moreover, the distribution of injuries can also indicate how injuries are managed in a species. Amputations at the trochanter by nestmates on ants with femur injuries have been reported in *Camponotus* species as an effective response to reduce infection risk (Frank et al., 2024; Fujimoto et al., 2025). If legs are amputated in a species, we would expect to see a decrease in the proportion of femur injuries and an increase in trochanter injuries. Our results indicate that *Lasius alienus* and *Temnothorax parvulus* showed a threefold higher rate of injuries at the trochanter than at the femur, whereas in most other species the opposite trend was observed. Interestingly, historical observations from the 1950s (Nachtwey, 1950) documented one amputation event in *L. alienus*, supporting the interpretation that *L. alienus* could use amputations as a behavioral response to injury and infection-related challenges.

## Conclusion

This study highlights that differences in injury rates and affected body regions among Central European ant species are shaped by feeding ecology characteristics such as diet and behaviour. Predatory species were more prone to leg injuries, while omnivorous species showed higher rates of antennal injuries, reflecting differences in foraging risks. Furthermore, the distribution of injuries could provide information on a species’ wound response strategies, such as leg amputations. Importantly, our findings demonstrate that pitfall traps are an effective tool for assessing injury rates across multiple species under natural conditions. Ultimately, our study highlights the ubiquity of injuries in ants, making it an important and measurable axis of selection that may have contributed to the emergence of wound care and other forms of social immunity.

## Author contribution

**Erik T. Frank** designed the study. **Alina Koeters** collected the data. **Melvin Opolka** analyzed the data. **Erik T. Frank** supervised and conceptualized the project. **Melvin K. Opolka** and **Alina Koeters** led the writing of the manuscript. All authors contributed critically to the drafts and gave final approval for publication.

## Acknowledgments

We thank Fabian S. Klimm for selecting and characterizing the study sites as part of a previous research project. We are also grateful to Christina Reith for her assistance during fieldwork, and to Franziska Engelhard for conducting the field assays involving the hand collection of *Lasius niger*.

## Funding

Erik T. Frank was supported by the DFG Emmy Noether Program #511474012 and the DFG grant #567282475.

## Conflict of interest

The authors declare that they have no known competing financial interests or personal relationships that could have influenced the work reported in this paper.

## Notes

### Competing Interest Statement

The authors have declared no competing interest.

