## Supplementary Information for "Feeding Ecology and Behavioral Adaptations Shape Injury Patterns in Central European Ants"

*Shared first authorship

**Short title:** Feeding Ecology Shapes Injury Patterns in Ants

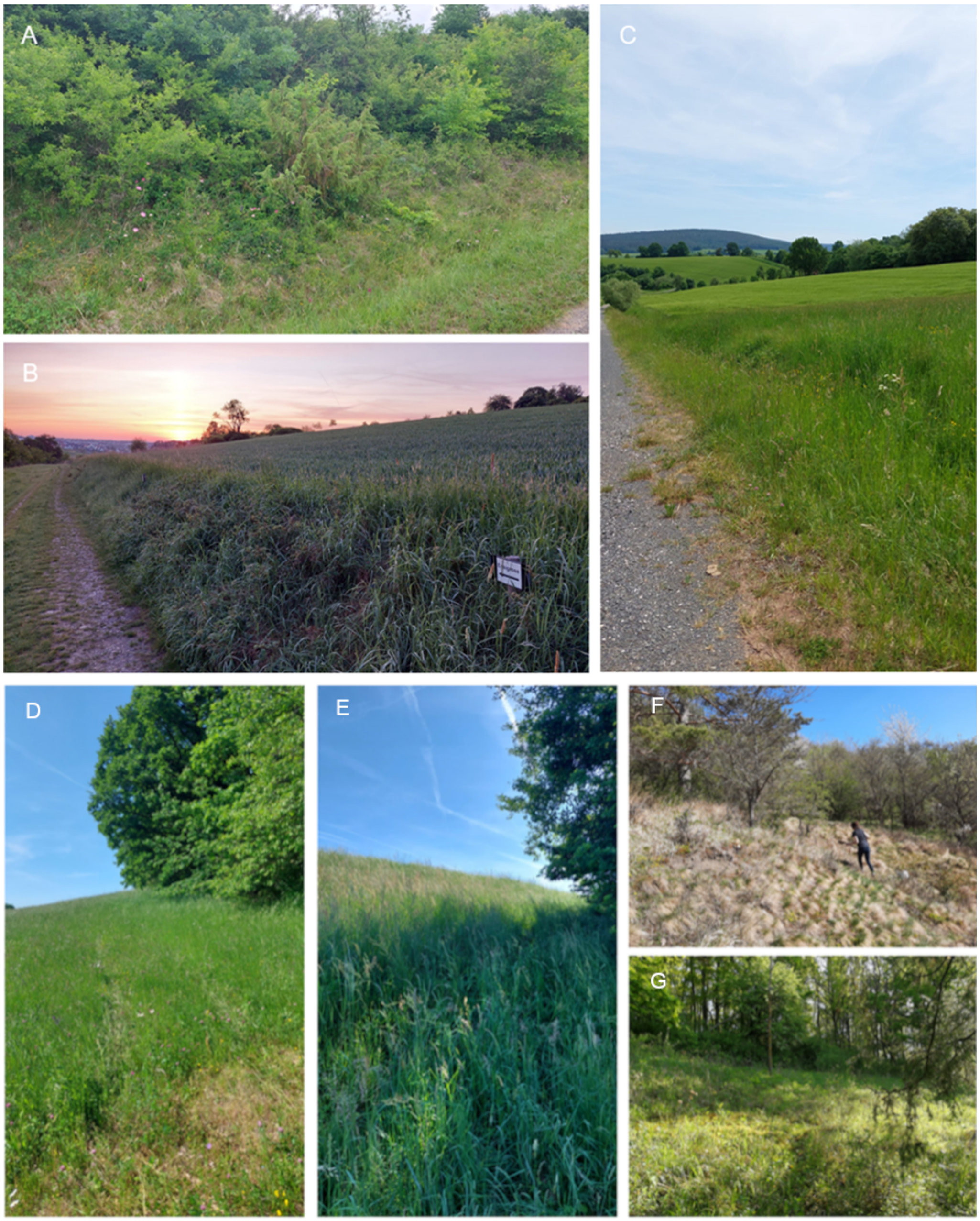


**Figure S1.** Locations with different habitats. (A) Gauaschach (slope), (B) Höchberg (wooded strip), (C) Gleusdorf (forest edge), (D) Krum (wooded strip), (E) Köslau (grassland), (F) Zimmerau (slope), (G) Unternesselbach (forest edge).

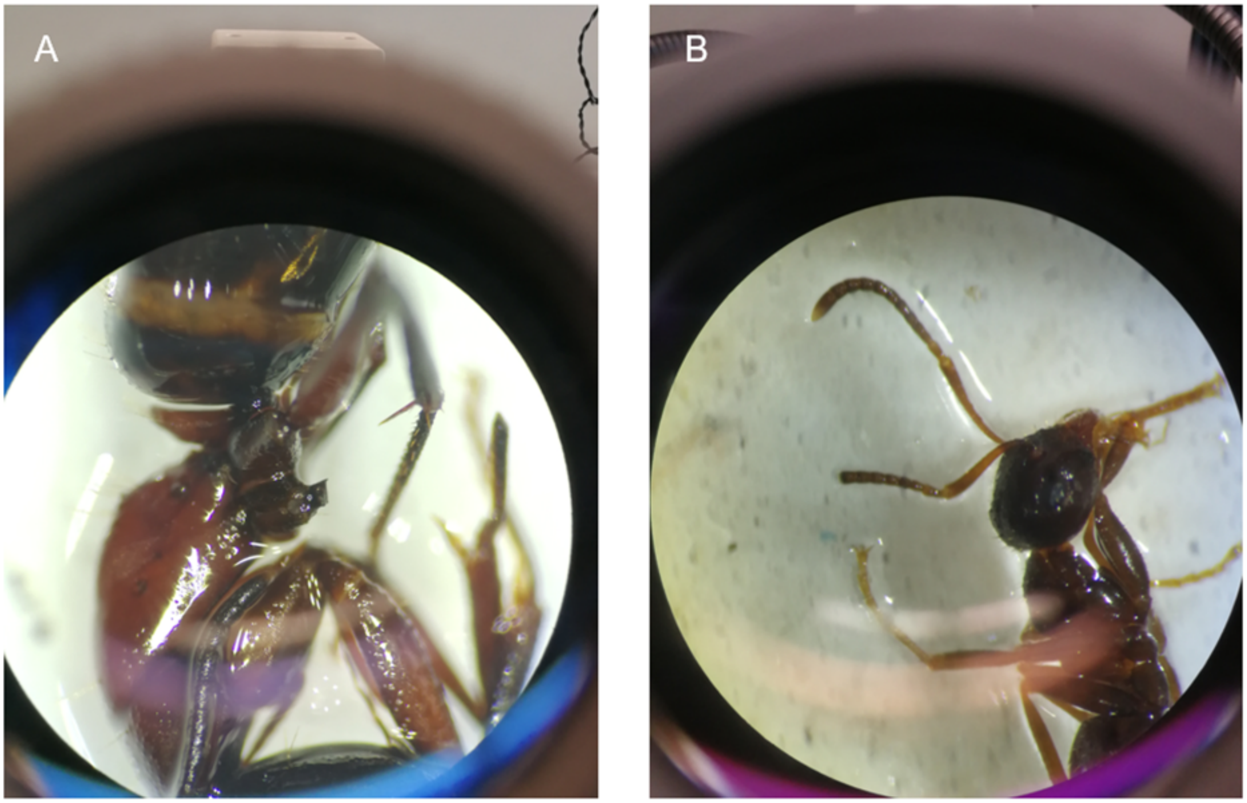


**Figure S2.** Photographs of injured ants. (A) *Camponotus ligniperda*, injury to the left middle trochanter, suggesting a possible amputation event conducted by nestmates as observed in *C. floridanus* (Frank et al. 2024); (B) *Lasius niger*, injury on the right funiculus.

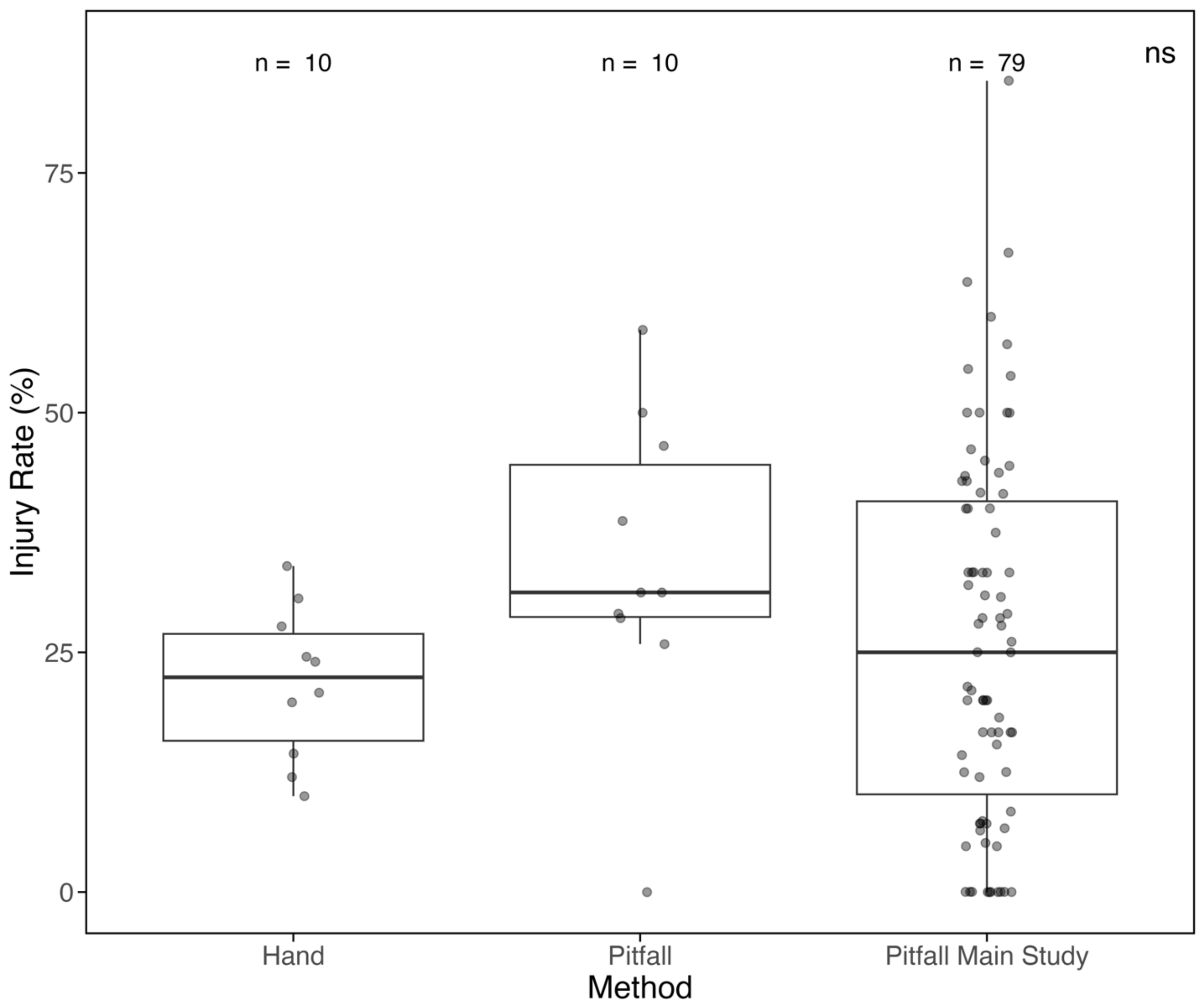


**Figure S3.** Comparison of injury rates in *Lasius niger* collected by hand versus pitfall traps; pitfall data from the main study are shown as an additional reference group. Sample sizes are indicated above each box. Each point represents one pitfall/colony. No significant difference was found between sampling methods or datasets (Wilcoxon signed-rank test, $p = 0.224$).

**Table S1.** Overview of the species. The table lists the subfamily, genus, and species, the number of individuals found in the samples, the number of injured individuals, the frequency of occurrence of the species across the 44 sites, and the species' feeding ecology.

| Subfamily | Genus | Species | Total number of captured individuals | Number of injured individuals | Frequency in sampling sites | Feeding ecology |
|---|---|---|---|---|---|---|
| Dolichoderinae | *Dolichoderus* | *quadripuncatus* | 2 | 1 | 1/44 | trophobiont |
| Dolichoderinae | *Tapinoma* | *erraticum* | 506 | 132 | 18/44 | omnivore |
| Formicinae | *Camponotus* | *herculeanus* | 3 | 1 | 2/44 | omnivore |
| Formicinae | *Camponotus* | *ligniperda* | 27 | 2 | 8/44 | omnivore |
| Formicinae | *Colobopsis* | *truncata* | 1 | 0 | 1/44 | omnivore |
| Formicinae | *Formica* | *cunicularia* | 29 | 5 | 8/44 | omnivore |
| Formicinae | *Formica* | *fusca* | 647 | 45 | 26/44 | omnivore |
| Formicinae | *Formica* | *polyctena* | 48 | 7 | 4/44 | predatory |
| Formicinae | *Formica* | *pratensis* | 2838 | 100 | 18/44 | predatory |
| Formicinae | *Formica* | *rufa* | 37 | 4 | 14/44 | predatory |
| Formicinae | *Formica* | *rufibarbis* | 79 | 11 | 15/44 | omnivore |
| Formicinae | *Formica* | *sanguinea* | 215 | 9 | 10/44 | predatory |
| Formicinae | *Lasius* | *alienus* | 579 | 148 | 20/44 | omnivore |
| Formicinae | *Lasius* | *brunneus* | 5 | 3 | 4/44 | trophobiont |
| Formicinae | *Lasius* | *flavus* | 19 | 6 | 9/44 | trophobiont |
| Formicinae | *Lasius* | *fuliginosus* | 55 | 6 | 12/44 | trophobiont |
| Formicinae | *Lasius* | *niger* | 1513 | 427 | 34/44 | omnivore |
| Formicinae | *Lasius* | *umbratus* | 2 | 1 | 1/44 | trophobiont |
| Myrmicinae | *Leptothorax* | *acervorum* | 1 | 0 | 1/44 | predatory |
| Myrmicinae | *Myrmecina* | *graminicola* | 43 | 0 | 12/44 | predatory |
| Myrmicinae | *Myrmica* | *rubra* | 312 | 16 | 18/44 | predatory |
| Myrmicinae | *Myrmica* | *ruginodis* | 1754 | 146 | 42/44 | predatory |
| Myrmicinae | *Myrmica* | *rugulosa* | 13 | 6 | 1/44 | omnivore |
| Myrmicinae | *Myrmica* | *sabuleti* | 5 | 0 | 2/44 | omnivore |
| Myrmicinae | *Myrmica* | *scabrinodis* | 42 | 8 | 6/44 | predatory |
| Myrmicinae | *Myrmica* | *schencki* | 15 | 3 | 4/44 | predatory |
| Myrmicinae | *Stenamma* | *debile* | 1 | 1 | 1/44 | predatory |
| Myrmicinae | *Temnothorax* | *albipennis* | 1 | 0 | 1/44 | omnivore |
| Myrmicinae | *Temnothorax* | *crassispinus* | 24 | 1 | 4/44 | omnivore |
| Myrmicinae | *Temnothorax* | *nylanderi* | 63 | 26 | 10/44 | omnivore |
| Myrmicinae | *Temnothorax* | *parvulus* | 586 | 52 | 23/44 | omnivore |
| Myrmicinae | *Temnothorax* | *unifasciatus* | 2 | 0 | 2/44 | omnivore |
| Myrmicinae | *Tetramorium* | *caespitum-impurum* | 7 | 2 | 3/44 | omnivore |

**Table S2.** Summary of all references used to classify ant species according to their respective feeding ecology.

| Ant species | Feeding ecology | Reference |
|---|---|---|
| *Dolichoderus quadripunctatus* | trophobiont | (Lebas et al., 2019) |
| *Tapinoma erraticum* | omnivore | (Lebas et al., 2019; Meudec & Lenoir, 1982) |
| *Camponotus herculeanus* | omnivore | (Ayre, 1963; Seifert, 2007) |
| *Camponotus ligniperda* | omnivore | (Seifert, 2007; Soares & Oliveira, 2021) |
| *Colobopsis truncata* | omnivore | (Lebas et al., 2019; Seifert, 2018) |
| *Formica cunicularia* | omnivore | (Lebas et al., 2019; Seifert, 2007) |
| *Formica fusca* | omnivore | (Lebas et al., 2019; Seifert, 2007) |
| *Formica polyctena* | predatory | (Véle & Modlinger, 2016) |
| *Formica pratensis* | predatory | (Novgorodova, 2015) |
| *Formica rufa* | predatory | (Lebas et al., 2019; Seifert, 2007) |
| *Formica rufibarbis* | omnivore | (Seifert, 2007) |
| *Formica sanguinea* | predatory | (Mori et al., 2000) |
| *Lasius alienus* | omnivore | (Akyürek et al., 2016; Kar et al., 2022) |
| *Lasius brunneus* | trophobiont | (Depa et al., 2022; Lebas et al., 2019) |
| *Lasius flavus* | trophobiont | (Parmentier & Wybouw, 2025) |
| *Lasius fuliginosus* | trophobiont | (Depa et al., 2022) |
| *Lasius niger* | omnivore | (Bulgarini et al., 2021; Lebas et al., 2019) |
| *Lasius umbratus* | trophobiont | (Depa et al., 2022) |
| *Leptothorax acervorum* | predatory | (Lebas et al., 2019) |
| *Myrmecina graminicola* | predatory | (Lebas et al., 2019) |
| *Myrmica rubra* | predatory | (Le Roux et al., 2002; Reznikova & Panteleeva, 2001) |
| *Myrmica ruginodis* | predatory | (Koptur & Lawton, 1988; O' Grady et al., 2010) |
| *Myrmica rugulosa* | omnivore | (Seifert, 2007) |
| *Myrmica sabuleti* | omnivore | (Lebas et al., 2019) |
| *Myrmica scabrinodis* | predatory | (O' Grady et al., 2010) |
| *Myrmica schencki* | predatory | (O' Grady et al., 2010) |
| *Stenamma debile* | predatory | (Lebas et al., 2019) |
| *Temnothorax albipennis* | omnivore | (Fokuhl et al., 2012; Seifert, 2018) |
| *Temnothorax crassispinus* | omnivore | (Fokuhl et al., 2012) |
| *Temnothorax nylanderi* | omnivore | (Fokuhl et al., 2012; Seifert, 2018) |
| *Temnothorax parvulus* | omnivore | (Fokuhl et al., 2012; Seifert, 2018) |
| *Temnothorax unifasciatus* | omnivore | (Lebas et al., 2019; Seifert, 2007) |
| *Tetramorium caespitum-impurum* | omnivore | (Lebas et al., 2019; Seifert, 2007) |

**Table S3.** Summary of three linear mixed-effects models (LMMs) assessing factors influencing injury rates in ants. All models include a random intercept for sampling location. Models differ in their fixed structure: (1) a full model with interaction between habitat and feeding ecology, (2) a model with feeding ecology only, and (3) a model with habitat only. AIC values are reported for model comparison.

| ***Response/Factor*** | ***Estimate*** | ***Std. error*** | ***t-value*** | ***p-value*** | ***AIC*** |
|---|---|---|---|---|---|
| *Injury_rate ~ habitat * feeding_ecology + (1\|location)* | | | | | 3064 |
| *Intercept* | 22.276 | 2.044 | 10.901 | **< 0.001*** ** | |
| *Grassland* | -0.903 | 3.717 | -0.243 | 0.81 | |
| *Slope* | -5.068 | 3.800 | -1.334 | 0.19 | |
| *Wooded strip* | -3.228 | 3.939 | -0.819 | 0.42 | |
| *Predatory* | -13.178 | 2.815 | -4.681 | **< 0.001*** ** | |
| *Grassland_Predatory* | -1.999 | 4.683 | -0.427 | 0.67 | |
| *Slope:Predatory* | 9.428 | 4.898 | 1.925 | 0.06 | |
| *Wooded Strip:Predatory* | 6.083 | 6.058 | 1.004 | 0.32 | |
| *Injury_rate ~ feeding_ecology + (1\|location)* | | | | | 3042.6 |
| *Intercept* | 20.682 | 1.298 | 15.939 | **< 0.001*** ** | |
| *Pedatory* | -11.177 | 1.810 | -6.174 | **< 0.001*** ** | |
| *Injury_rate ~ habitat + (1\|location)* | | | | | 3113 |
| *Intercept* | 17.4851 | 1.9838 | 8.814 | **< 0.001*** ** | |
| *Grassland* | -3.5326 | 3.4265 | -1.031 | 0.31 | |
| *Slope* | -1.977 | 3.5482 | -0.557 | 0.58 | |
| *Wooded strip* | -0.7529 | 3.7908 | -0.199 | 0.84 | |

**Table S4.** Summary of fixed effects from the linear mixed-effects model assessing injury rates across ant body parts and feeding ecologies.

| ***Response/Factor*** | **Estimate** | **Std. error** | **t-value** | **p-value** |
|---|---|---|---|---|
| *Injury_rate ~ feeding_ecology *body_part (1\|sampleID)* | | | | |
| *Intercept* | 21.17 | 0.72 | 29.40 | **< 0.001***** |
| *Predatory* | -16.71 | 1.12 | -14-94 | **< 0.001***** |
| *Front Legs* | -18.23 | 0.97 | -18.84 | **< 0.001***** |
| *Middle Legs* | -19.91 | 0.97 | -20.57 | **< 0.001***** |
| *Hind Legs* | -20.1 | 0.97 | -20.74 | **< 0.001***** |
| *Predatory:Front Legs* | 15.77 | 1.50 | 10.49 | **< 0.001***** |
| *Predatory:Middle Legs* | 17.02 | 1.50 | 11.33 | **< 0.001***** |
| *Predatory:Hind Legs* | 18.32 | 1.50 | 12.19 | **< 0.001***** |
